# Influence of genetic polymorphism on transcriptional enhancer activity in the malaria vector *Anopheles coluzzii*

**DOI:** 10.1101/659144

**Authors:** Luisa Nardini, Inge Holm, Adrien Pain, Emmanuel Bischoff, Daryl M Gohl, Soumanaba Zongo, Wamdaogo M. Guelbeogo, N’Fale Sagnon, Kenneth D Vernick, Michelle M Riehle

## Abstract

Enhancers are cis-regulatory elements that control most of the developmental and spatial gene expression in eukaryotes. Genetic variation of enhancer sequences is known to influence phenotypes, but the effect of enhancer variation upon enhancer functional activity and downstream phenotypes has barely been examined in any species. In the African malaria vector, *Anopheles coluzzii*, we identified a pilot set of candidate enhancers in the proximity of genes relevant for immunity, insecticide resistance, and development. The candidate enhancers were functionally validated using luciferase reporter assays, and their activity was found to be essentially independent of their physical orientation, a typical property of enhancers. All of the enhancer intervals segregated genetically polymorphic alleles, which displayed significantly different levels of functional activity, and inactive null alleles were also observed. Deletion mutagenesis and functional testing revealed a modular structure of positive and negative regulatory elements within the tested enhancers. The enhancer alleles carry genetic polymorphisms that also segregate in wild *A. coluzzii* populations in West Africa, indicating that enhancer variants with likely phenotypic consequences are frequent in nature. These results demonstrate the feasibility of screening for naturally polymorphic *A. coluzzii* enhancers that underlie important aspects of malaria transmission and vector biology.

## INTRODUCTION

Enhancers are short cis-acting regulatory elements in noncoding DNA that amplify transcriptional levels of target genes by tens to hundreds fold over the basal level of core promoter elements at the transcription start site. Core promoters are located proximal to the transcription start site (TSS) and facilitate the binding of RNA polymerase and the initiation of transcription ^1–3^. Enhancers control transcriptional activity of target genes through their interaction with activators and the promoter and are, in turn, responsible for most regulated gene expression in the transcriptome. The precise mechanisms of enhancer action is unknown and is an area of active study ^4–6^. Enhancers can function at a distance from target genes and independent of their physical orientation in the chromosome ^7^. The identities of enhancers and some of their interacting protein factors that lead to their regulatory function have been described in well-studied model organisms, but enhancers cannot be reliably predicted by sequence-based algorithms, and thus must be detected directly by functional activity using reporter assays, or indirectly inferred using methods to detect open or modified chromatin properties.

Sequence polymorphism within enhancers has been associated with phenotypic differences, including predisposition to disease, as observed in diverse organisms ^5, 8–11^. Most of the significant variants mapped in human genome-wide association studies (GWAS) are noncoding ^12^, and at least 60-70% of significantly associated human GWAS single-nucleotide point mutations (SNPs) lie within functional enhancers ^8–13^. In cancer studies, the majority of tumor-driving mutational changes are also thought to be in noncoding regulatory elements, especially enhancers ^14^.

Genetic variation in enhancers can occur as SNPs, insertions and deletions (indels), and as copy number variants ^15–17^. Enhancer variation among individuals can underlie both Mendelian and complex traits ^7–18, 19^. At the population level, positively-selected variation in enhancers controlling key pathways likely plays an important role in differentiation and evolution ^20, 21^. Indeed, some of the fastest-evolving parts of the human genome as compared to other primates are functional embryonic enhancers related to central nervous system development ^22^. In another example, the vertebrate ZRS enhancer influencing limb development displays strong conservation across a range of vertebrates, although in advanced snakes where skeletal limb structures are absent, mutations render the enhancer inactive ^23^. Finally, stickleback fish display development of lateral pelvic spines in some populations, which may be protective under certain predation regimes but may be an adaptive liability in others, and spines have been independently lost in different populations. Sequence differences in the Pitx1 enhancer among spined and spineless populations correlate with the phenotype, and enhancer swaps restore or abolish spine development ^24^. Work in Drosophila has shown that enhancer variation effects many aspects of morphology including for example pigmentation ^25^, melanism ^26^ and larval tricome pattern and number ^27^. Numerous Gal4 based enhancer trapping lines exist for Anopheles stephensi largely as a tool for the study mosquito functional genomics and optimizing methods to drive transgene expression ^28^. Recent work in the African malaria vector Anopheles funestus, highlights the profound effects of indel variation at a cis regulatory element in the enhancer/promoter region of a cytochrome P450 and its association with insecticide resistance ^29^. Thus, relatively simple evolved sequence variation in enhancers can produce large phenotypic shifts in the organism ^30, 31^. Despite these examples, the effect of genetic variation on enhancers has barely been examined in any species, and to our knowledge, the functional 4 effect of variation on enhancer activity has not been systematically surveyed in any organism.

Enhancer positional effects on gene expression are known to influence phenotype and can lead to disease states. Viruses and mobile elements carrying an enhancer can place surrounding genes under a different regulatory regime, as well as chromosome translocations such as the human ‘Philadelphia chromosome’ and others that promote <u>chronic myeloid leukemia</u>. In genetic vector control strategies, it could be important to know the locations of enhancers that could cause unpredicted effects on transgene expression.

Enhancer discovery in mosquitoes is limited to a few previous studies using indirect methods based on detection of chromatin properties to infer enhancer locations ^32–34^. However, almost nothing is known about mosquito enhancer genetic variation and influence on vector phenotypes, and thus it is necessary to first develop the baseline criteria to distinguish and study enhancers in *A. coluzzii*. To this end, here we analyze a pilot set of candidate enhancers in close proximity to a small number of genes of interest in order to benchmark the parameters needed for reliable enhancer discovery, validation, and determination of polymorphism effects in *Anopheles*. We validate the pilot candidates as functional enhancers using luciferase reporter assays, and measure the effects of genetic polymorphism on enhancer activity. The results of the current report are a first step towards developing a comprehensive genome-wide catalog of *Anopheles* enhancers, and biological studies to characterize enhancer function in vector biology.

## RESULTS

### Candidate enhancer selection

The standard approach for enhancer detection is by functional testing using luciferase reporter assays that directly measure enhancer activity, or by indirect methods such as ChIP-seq, which can infer the presence of a subset of enhancers by correlation with chromatin features. Here, we implemented for the first time in *Anopheles* a screen (Self-Transcribing Active Regulatory Region sequencing, STARR-seq) that detects enhancers directly by a functional reporter assay analogous to the luciferase reporter assay, but with the readout of enhancer-dependent RNA transcript output measured as sequenced cDNA, rather than by luciferase light output ^35^.

In order to identify candidate *A. coluzzii* enhancers, we examined our generated sequence data from the functional screen in the vicinity of six selected genes using the Integrative Genomics Viewer (IGV) ^36^. Candidate enhancers were identified by using IGV to manually search near the selected genes to detect intervals where coverage of the cDNA sequence track, indicative of enhancer activity in *A. coluzzii* 4a3A cells (Figure 1, solid lines) was visibly greater than the coverage of the plasmid sequence track, which is the internal baseline control indicating background levels of the plasmid in 4a3A cells (Figure 1, dotted lines). The target genes selected are involved in vector immunity: Krueppel-Like Factor 6/7 (KLF, AGAP007038), Leucine-Rich Immune protein (LRIM1, AGAP006348); insecticide resistance: Acetylcholinesterase (ACE1, AGAP001356), GABA-gated chloride channel subunit (Rdl, AGAP006028); and developmental biology: LIM homeobox protein 2/9, ortholog of Drosophila apterous FBgn0267978 (AP, AGAP008980), and Ovo, AGAP000114 (Table 1). The regulatory regions and enhancers of the latter two genes, apterous and Ovo, have been well characterized for the Drosophila orthologs ^37–39^. In addition, a negative control interval was chosen as a size-matched interval located within intron 1-2 of the gene, homeobox protein distal-less (DLX, AGAP007058) that has no visible divergence of cDNA and plasmid sequence tracks by IGV examination, and thus no predicted enhancer function. The candidate enhancer intervals are named according to the most proximal coding sequence.

**Figure 1.**
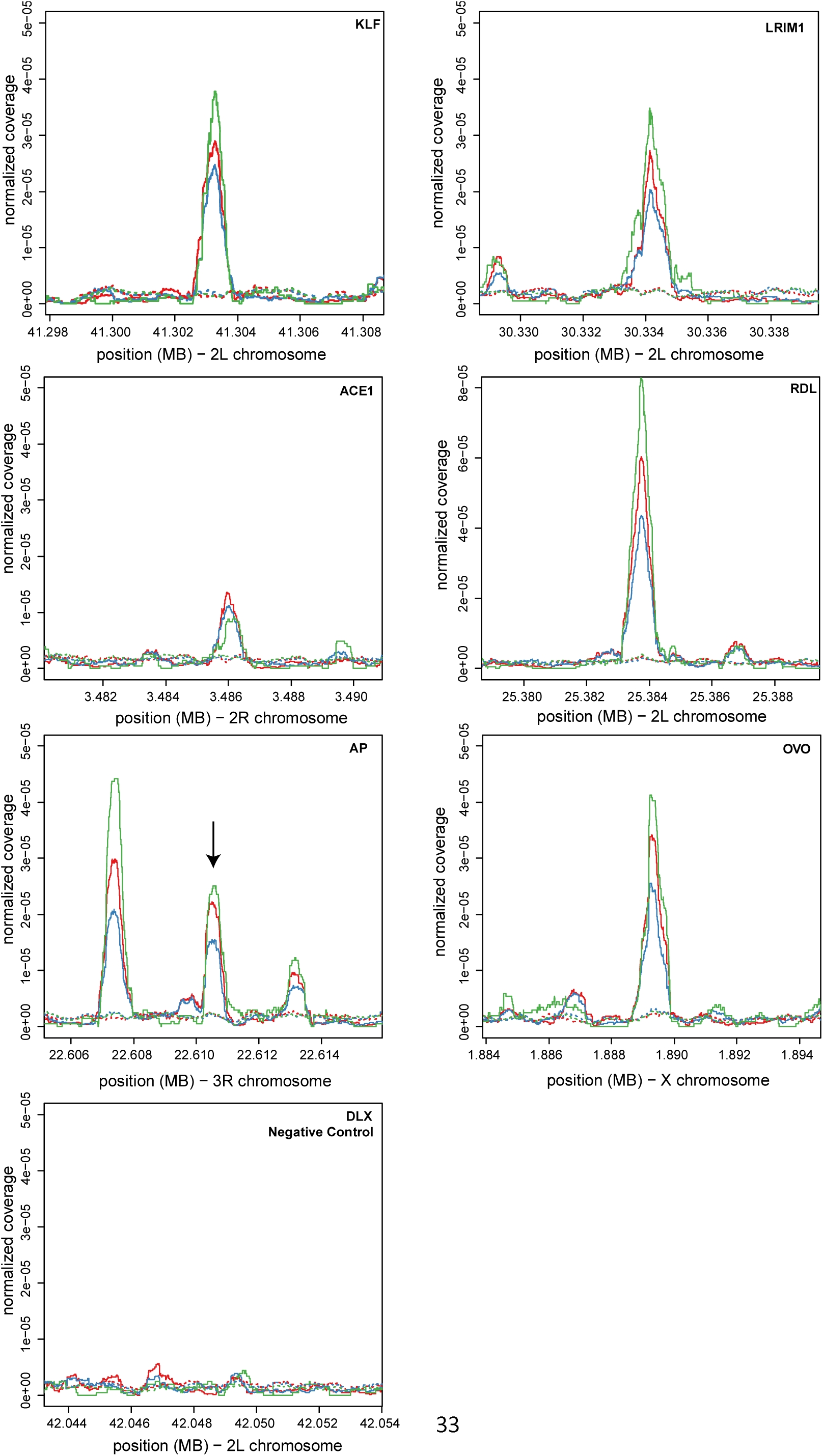
Detection of *Anopheles coluzzii* candidate enhancers. Sequence data near six *Anopheles coluzzii* genes were examined using the Integrative Genomics Viewer (IGV) ^36^ to screen for candidate enhancers, identified visually where coverage of the cDNA sequence track (solid lines) was greater than the baseline coverage of the plasmid sequence track (dotted lines). The cDNA sequence track is analogous to light output from luciferase reporter assays, but where the readout is enhancer-dependent RNA transcript output in *A. coluzzii* 4a3A cells, measured as sequenced cDNA, rather than by luciferase light output. Line color (green, red, blue) represents three biological replicates. The enhancers are named by the most proximal genes: Krueppel-Like Factor 6/7 (KLF, AGAP007038), Leucine-Rich Immune protein (LRIM1, AGAP006348), Acetylcholinesterase (ACE1, AGAP001356), GABA-gated chloride channel subunit (Rdl, AGAP006028), LIM homeobox protein 2/9, ortholog of Drosophila apterous FBgn0267978 (AP, AGAP008980), and Ovo, AGAP000114 (Table 1). A negative control interval within intron 1-2 of distal-less (DLX, AGAP007058) was chosen because it lacks visible divergence of cDNA and plasmid sequence tracks. Graphs display cDNA and plasmid tracks in 10 kb windows centered on the candidate enhancers. Only one candidate enhancer is seen in all windows except AP, where the central peak (arrow) was used. X-axis indicates genomic coordinates in the PEST reference genome, y-axis indicates normalized sequence depth corrected for overall plasmid depth observed in the IGV display.

### Functional validation of enhancer activity

The predicted candidate enhancers predicted in Figure 1 were functionally tested to validate the IGV-based predictions. The standard test for enhancer activity is by cloning the candidate in a plasmid carrying a basal promoter and a luciferase reporter gene in an episomal assay. An active enhancer will augment the rate of transcription from the basal promoter, thus elevating the expression level of the luciferase gene. Luciferase expression is measured by adding luciferase substrate to cell extract and detecting light output as relative light units (RLU). Enhancer activity, if any, is measured as increased luciferase activation above background.

Candidate amplicons from A. *coluzzii* (Table 1) were cloned into the firefly luciferase reporter vector pGL-Gateway-DSCP, and co-transfected into 4a3A cells with the renilla luciferase control vector pRL-ubi-63E. Firefly luciferase RLU measurements were corrected using the renilla luciferase internal control values in the same well, and firefly/renilla RLU for the experimental insert were statistically compared to the firefly/renilla mean value for the DLX negative control, defined as the background level. At least one clone of each candidate enhancer displayed luciferase activity levels above background (p<0.005), with activity across candidates that varied from 2-fold to more than 20-fold over background (Figure 2). These results indicate that the IGV-based predictions were accurate for all six predicted candidates, and thus validate these genomic intervals as functional *A. coluzzii* enhancers in the cell culture model. Further work will be required to determine enhancer activity in the native chromosomal context. The current information provides the first benchmark criteria that can be used to develop the definitions and methods for subsequent algorithmic genome-wide detection of *A. coluzzii* enhancers.

**Figure 2.**
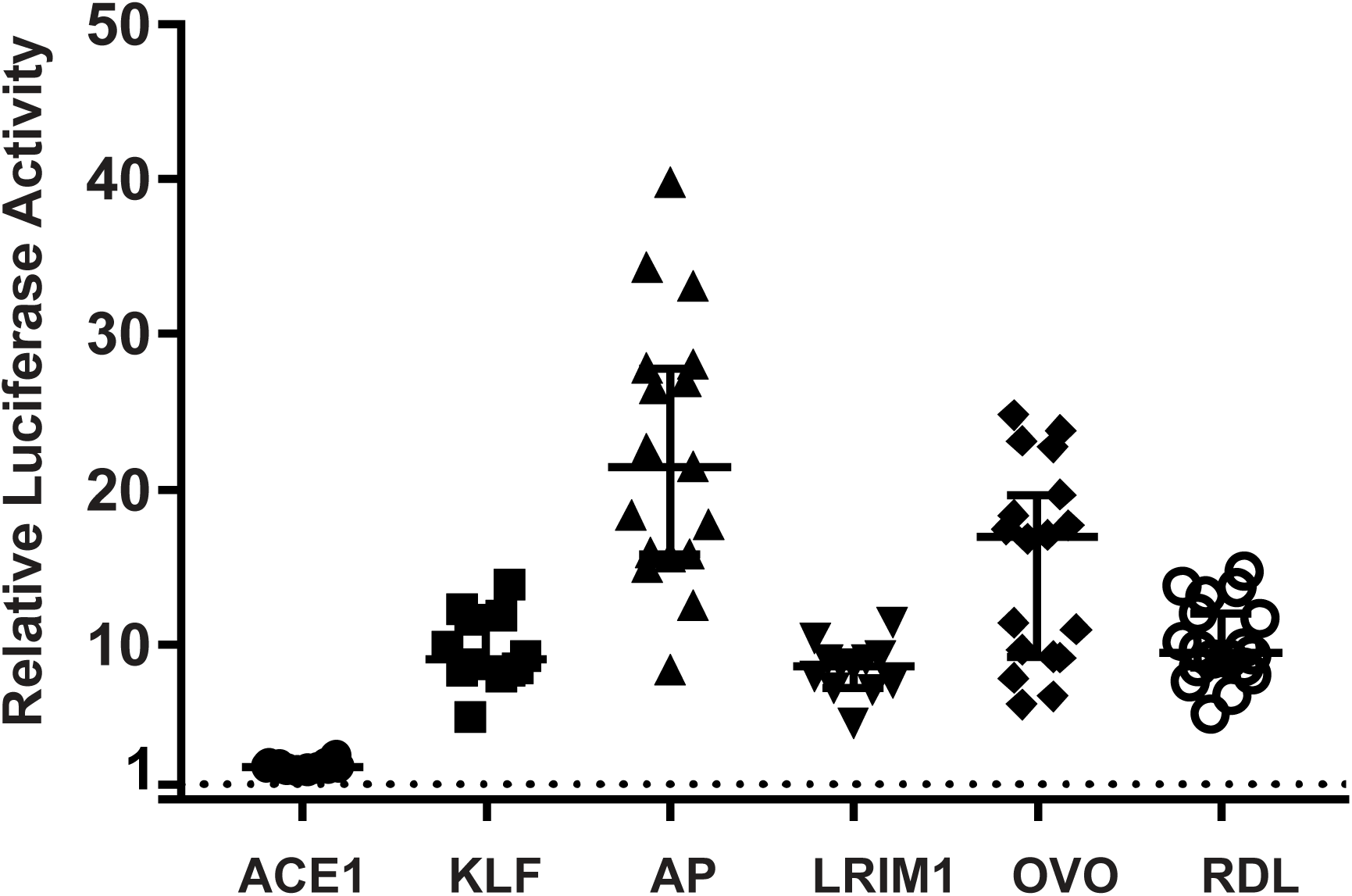
Candidate *Anopheles coluzzii* enhancers augment expression of a luciferase reporter. The six candidate enhancer intervals from Figure 1 were amplified from *Anopheles coluzzii* mosquitoes and cloned into the pGL-Gateway-DSCP plasmid carrying a basal core promoter and firefly luciferase reporter gene. The cloned candidates were tested for influence upon luciferase expression using a dual luciferase assay system to quantify luciferase activity above background, defined by the DLX negative control (horizontal dotted line). Each of the six tested candidates displayed normalized luciferase activity significantly above background (p<0.005), thus validating the candidates as functional *A. coluzzii* enhancers. Each point represents an individual replicate measure of luciferase activity for the tested candidate. Bars indicate the median and 95% confidence intervals. X-axis indicates the name of the candidate enhancer according to the nearest gene (Table 1), y-axis indicates the relative luciferase activity for each measurement, expressed as firefly luciferase corrected to the renilla luciferase internal control value, and normalized for the value of the DLX negative control (see Methods).

**Table 1:** Physical location of enhancer intervals and proximal annotated gene. Enhancer interval coordinates are based on the locations of PCR primers given in Supplementary Table S2. Coordinates from the PEST AgamP4 genome assembly.

| Proximal gene |  | Enhancer Interval | Proximal Gene Coordinates |
| --- | --- | --- | --- |
| AP | (AGAP008980) | 3R:22609939-22611138 | 3R:22543990-22609635 |
| OVO | (AGAP000114) | X:1888505-1890055 | X:1852650-1884326 |
| KLF | (AGAP007038) | 2L:41302647-41303886 | 2L:41287202-41308450 |
| LRIM1 | (AGAP006348) | 2L:30333431-30334787 | 2L:30329656-30331296 |
| ACE1 | (AGAP001356) | 2R:3485436-3486583 | 2R:3483099-3497400 |
| RDL | (AGAP006028) | 2L:25382828-25384253 | 2L:25363652-25434556 |

### Screening for polymorphic alleles of validated enhancer intervals

Having confirmed that all six predicted candidates are functional enhancers, we next wished to identify genetically variable alleles for each enhancer interval and measure their luciferase activity. For this, alleles of the enhancer intervals were amplified and sequenced from *A. coluzzii* colonies initiated from the populations in Cameroon, Mali or Burkina Faso. For each of the six enhancer intervals, at least two distinct genetic variants were chosen for tests of enhancer activity. The goal was to identify and test a range of genetic variants, not the mosquito colonies. Therefore, a given colony may or may not be represented for a given enhancer, depending on the variation it segregates. The enhancer interval alleles were cloned and sequenced, and neighbor joining (N-J) trees depict the evolutionary relatedness and degree of sequence difference of the alleles (Figure 3). Complete sequences for all tested enhancer interval alleles are presented in Supplementary File S1.

**Figure 3.**
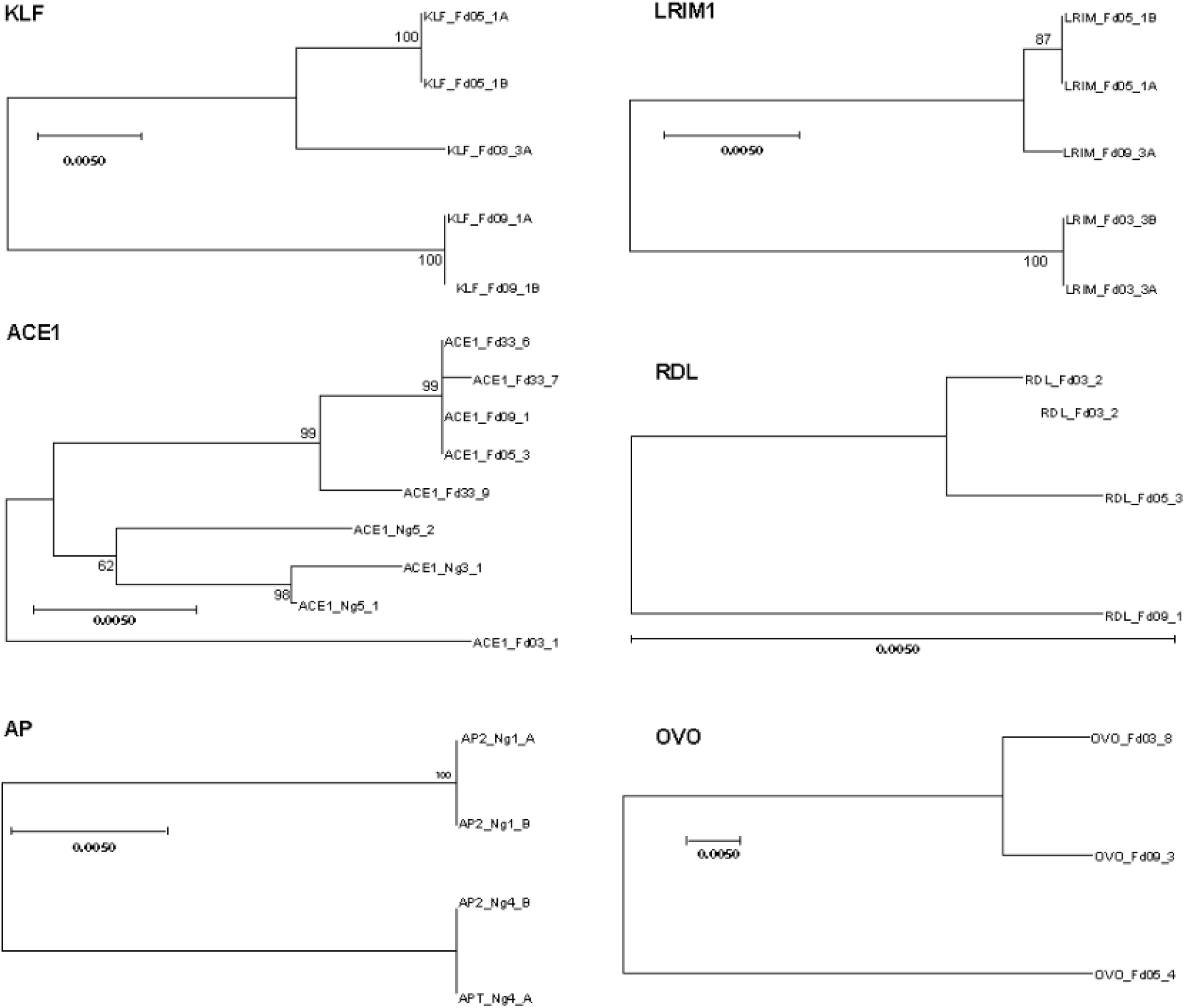
Phylogenetic comparison of enhancer allele genetic variation. Enhancer allelic variants were cloned and sequenced from *Anopheles coluzzii* colonies. Each sequenced clone represents a chromosomal haplotype. For each clone, individual sequences were aligned using MUSCLE and Mega was used to construct neighbor joining (N-J) trees for complete sequences for all haplotypes for each enhancer. Trees depict the degree of genetic similarity among alleles, and phylogenetic scale bars represent 0.5 nucleotide substitutions per site. The scale bar for the Rdl tree is long (pairwise distance 0.008 between alleles Fd03_#2 and Fd09_#1), indicating that the Rdl alleles segregate relatively little variation, while the Ovo tree scale bar is short (pairwise distance 0.0445 between alleles Fd03_#8 and Fd05_#4), indicating more than 5-fold greater genetic diversity among Ovo alleles as compared to Rdl. Alignments for complete sequences of alleles are presented in Supplementary File S1.

### Genetic alleles of validated enhancer intervals display distinct enhancer activity

Luciferase activity was measured for all alleles to determine the effect of genetic variation on differences in functional enhancer activity. For five of the six enhancers, alleles displayed significantly different levels of enhancer activity (Figure 4). For a given enhancer, alleles with 8 the greatest difference in activity tended to be the most genetically different from each other (see also Figure 3). For example, the alleles of the KLF interval cloned from colonies Fd05 and Fd03 are the most closely related genetically, and these also do not display a difference in luciferase activity as compared to the allele from colony Fd09. For two enhancer intervals (LRIM1 and ACE1), at least one genetic variant displayed activity levels that were not significantly different from background, which effectively represents a naturally occurring functionally inactive null enhancer allele. Genetic variation segregating at the enhancer of Ovo did not display a significant influence on luciferase activity, and the Ovo enhancer appears to display the consistently highest luciferase activity over all alleles tested for any of the six enhancer intervals. These results indicate that genetic alleles of validated enhancer intervals can display significantly different levels of functional activity. It is reasonable to expect that large observed differences in enhancer activity will be translated into phenotypic differences in the organism, related to the functions of the target genes that are regulated by the polymorphic enhancer alleles. This prediction will need to be tested in manipulative experiments.

**Figure 4.**
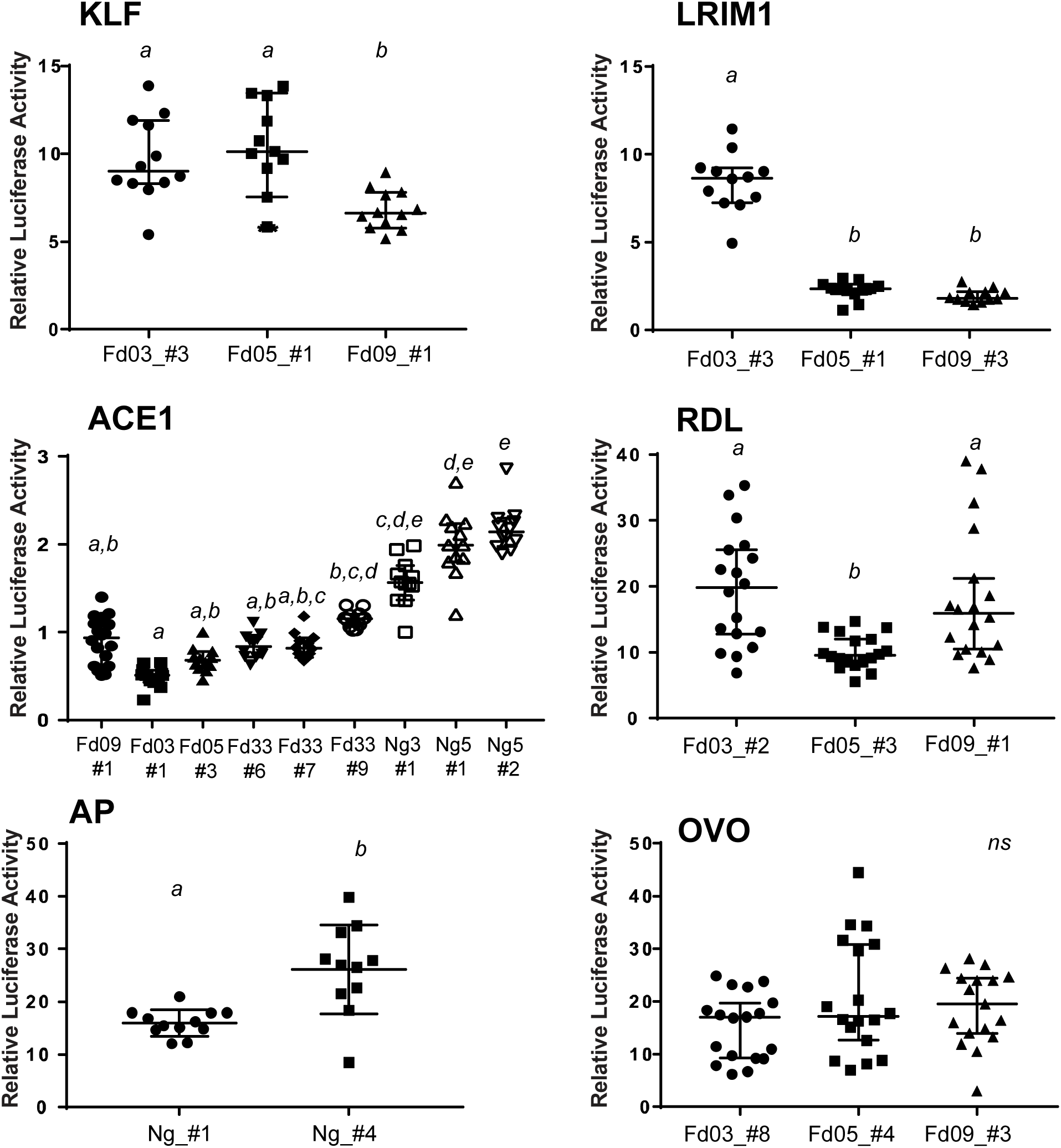
Genetic variation influences enhancer functional activity. To test the functional effect of genetic variation within enhancer sequences, the enhancer alleles shown in Figure 3 were cloned into luciferase reporter plasmid pGL-Gateway-DSCP, and luciferase activity was measured. Statistically significant differences in luciferase activity as determined using a non-parametric ANOVA are indicated by letters, samples labelled with different letters are significantly different from each other and samples with the same letter are not significantly different (thus samples labeled a,b are not statistically different from samples labelled either a or b). Bars indicate the median and 95% confidence intervals, n=12 for all tests. X-axis labels indicate colony origin (Ng, Ngousso, other colony names as given) and allele name, y-axis indicates the relative luciferase activity for each measurement determined as in Figure 2.

### Enhancer activity is essentially independent of physical orientation

Enhancers tend to function independently of their physical orientation in the genome, which is testable when the candidate is cloned in a luciferase reporter plasmid. For three of the above validated enhancers, we recloned two alleles in both orientations in the reporter and measured luciferase activity. For the KLF and AP enhancers, there was no detectable effect of orientation (Figure 5). The LRIM1 enhancer displayed a weakly significant effect of orientation for allele Fd05_#1 (p=0.042), although for both orientations of the LRIM1 enhancer the absolute activity values were lower than the other enhancers tested (indicated 9 by y-axis values in Figure 5), and thus the weak orientation difference for this one weak allele is not robustly supported. Thus, most of the enhancer alleles tested displayed function independent of their physical orientation with respect to the basal promoter.

**Figure 5.**
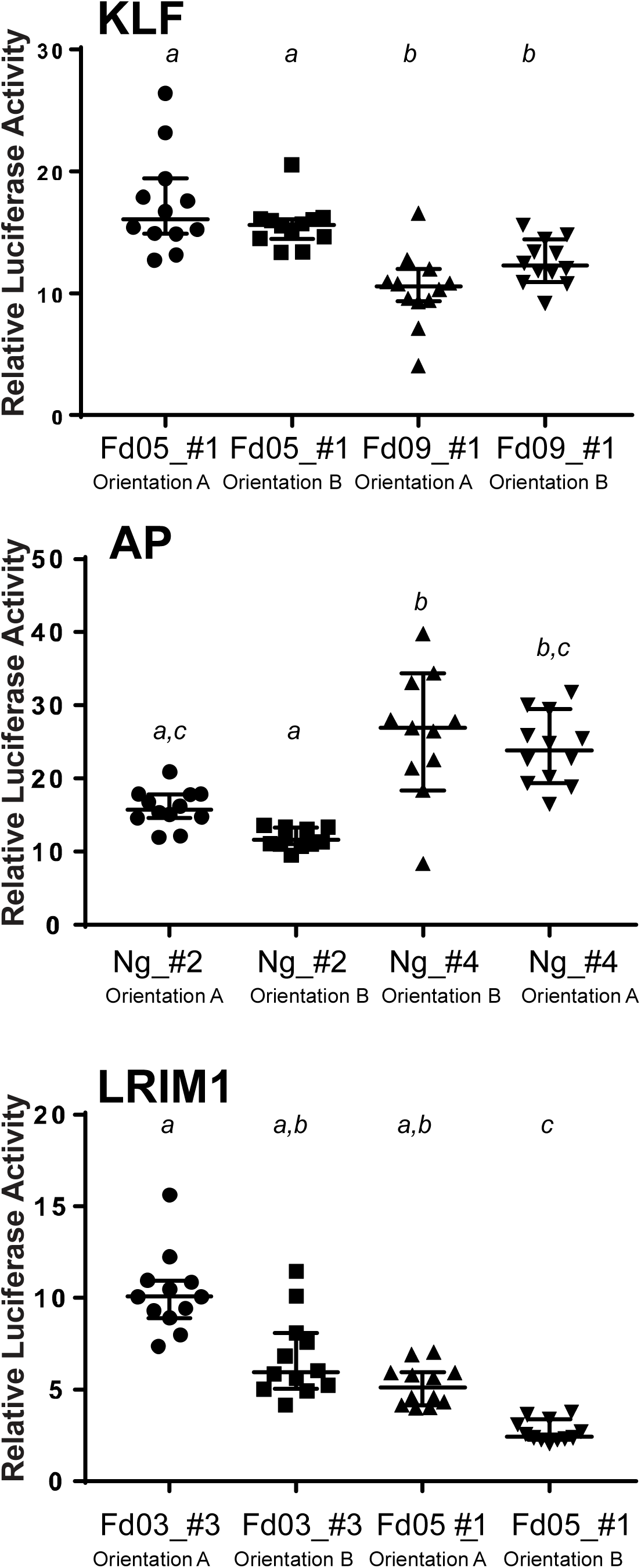
Enhancer activity is essentially independent of orientation. The influence of physical orientation of the enhancer within the luciferase reporter plasmid pGL-Gateway-DSCP was tested by cloning three enhancers, KLF, AP and LRIM1, in both orientations in the plasmid, and luciferase activity was measured. KLF and AP enhancers displayed no detectable effect of orientation on luciferase activity, while LRIM1 displayed a slightly significant difference (p=0.042) in luciferase activity for the allele Fd05_#1. Statistical differences indicated by letters as in Figure 4, error bars as in Figure 4, n=12 for all tests. X-axis indicates the name of the enhancer allele tested and the enhancer insert orientation (arbitrarily defined as A and B), n indicates the number of wells measured, y-axis indicates the relative luciferase activity for each measurement as in Figure 2.

### Enhancer deletion mutagenesis reveals a modular structure of positive and negative regulatory elements

To resolve the minimal portion of the enhancer interval that carries the enhancer function, we carried out deletion mutagenesis for two different genetic alleles of the LRIM1 enhancer interval, one allele with high enhancer activity and the other low. The LRIM1 enhancer was used for proof of principle, and was chosen because it had alleles with distinct activity levels, and the genetic variation was spread across the enhancer interval. The deletion derivatives carried 50% or 25% of the length of the initial enhancer interval, reduced equally from both ends. We tested the deletion clones for luciferase activity, along with the original undeleted enhancer (Figure 6A). Surprisingly, for LRIM1 allele Fd03_#3, the 50% construct displayed the highest luciferase activity, greater than either 100% or 25% constructs. This indicates that the integral 100% Fd03_#3 allele carries negative regulators of enhancer function, which were deleted in the 50% derivative to yield a derivative with elevated enhancer activity. The 25% derivative of allele Fd03_#3 displays significantly lower activity than the 50% derivative, suggesting that positive regulators of enhancer function are located outside the 25% derivative, but within the 50% derivative.

**Figure 6.**
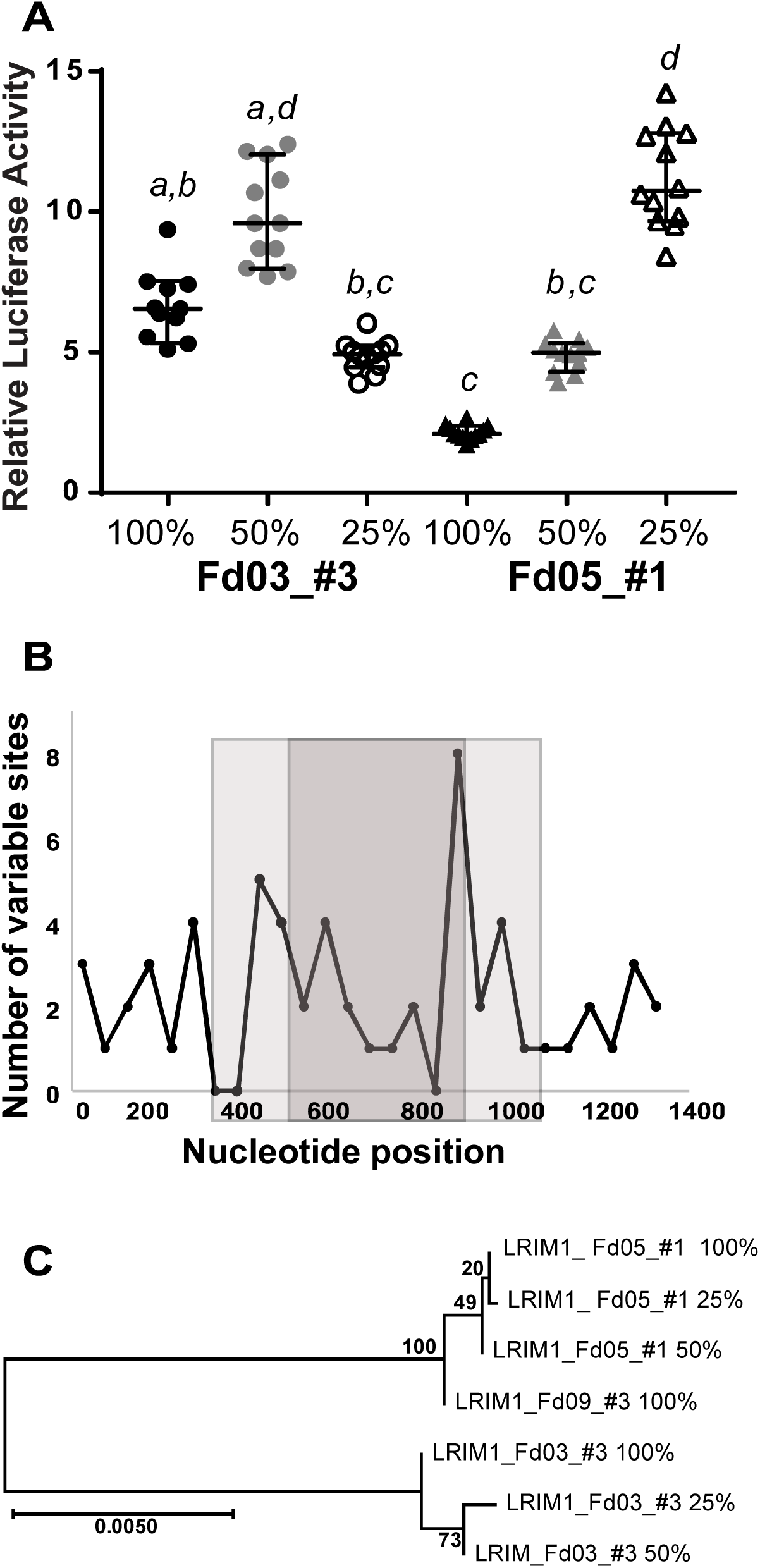
Enhancer deletion mutagenesis reveals positive and negative regulatory elements. Deletion mutagenesis was carried out for two alleles of the LRIM1 enhancer, the high-activity allele Fd03_#3 and low-activity allele Fd05_#1 (Figures 4 and 5). The integral enhancer alleles (100%) were each deleted for one-quarter of their length from both termini (50% derivative), and one-quarter length again (25% derivative). **A.** Deletion derivatives were tested for luciferase activity, along with the original integral alleles. Statistical differences indicated by letters as in Figure 4, error bars as in Figure 4, n=12 for all tests. X-axis indicates allele name and deletion derivatives, y-axis indicates the relative luciferase activity for each measurement as in Figure 2. Enhancer activity is not directly correlated with sequence length, and enhancer alleles are structured from distinct combinations of positive and negative regulators of transcription. **B.** Plot of the number of variant nucleotide positions between the Fd03_#3 and Fd05_#1 alleles along the length of the enhancer sequence. Variant sites are counted within a 50 bp non-overlapping window and plotted at the midpoint of the window. The light gray shading indicates the extent of the 50% length derivatives and the dark gray shading the 25% derivatives. X-axis indicates nucleotide position in derivatives, y-axis indicates number of variable sites between the Fd03_#3 and Fd05_#1 alleles in 50 bp windows. There were a total of 60 variable sites between Fd03_#3 and Fd05_#1 alleles in the 100% integral enhancer, 37 variable sites in the 50% derivatives and 22 sites in the 25% derivatives. **C.** Neighbor-joining tree depicting sequence relatedness between the integral 100% enhancer and the 50% and 25% derivatives for LRIM1 Fd03_#3 and Fd05_#1 alleles. The Fd09_#3 allele is included as an outgroup. Scale bar description as in Figure 3.

Deletion derivatives of LRIM1 allele Fd05_#1 display a pattern distinct from the Fd03_#3 allele. For Fd05_#1, each incrementally smaller derivative was more active. This result was also surprising, because it indicates that a highly active core enhancer element within the 10 smallest 25% derivative is attenuated by negative regulators that are progressively removed from 100% to 50% in length, and again from a 50% to 25% length interval. The deletion results indicate that enhancer activity is not directly correlated with sequence length, that there is a complex structure of functional elements and modifiers within the enhancer interval, and that different alleles of the same enhancer are comprised of distinct combinations of modular regulators that differentially influence transcription.

The density of variable sites between Fd03_#3 and Fd05_#1 varies across the interval, such that there were 60 variable nucleotide sites in the integral 100% length alleles, 37 variable sites in the 50% derivatives and 22 sites in the 25% derivatives (Figure 6B). Finally, it is notable that the 25% derivative for allele Fd05_#1 displays activity levels indistinguishable from the Fd03_#3 50% derivative (p=0.99), even though they are no more genetically similar than the integral 100% enhancer sequences for both alleles (Figure 6C). This result highlights the relative independence of enhancer functional level from primary sequence patterns, unlike the fundamental dependence of protein coding gene function on the amino acid primary sequence code, and the consequent requirement for identification of enhancers by detecting functional activity. Testing of a large panel and manipulative experiments would be required to identify consistent patterns of enhancer modular organization.

### Enhancer alleles segregate in the natural *Anopheles coluzzii* population

To confirm that the genetic variation observed in the enhancer alleles was natural and not an artifact of laboratory colonies, we compared sequence data for the six enhancer intervals to genetic variation observed in wild *A. coluzzii* from whole genome sequence of the *Anopheles* gambiae 1000 (Ag1000) Genomes Consortium ^40^. The comparison indicates that 11 genetic variation is shared between the cloned *A. coluzzii* colony haplotypes used in luciferase assays and the natural population (Figure 7). Representative short sequence alignments are shown, and full-length alignments with larger numbers of wild mosquitoes are presented in Supplementary File S2. Alignments are presented rather than phylogenetic trees because the wild Ag1000 sequences were called for SNPs but not indels. Therefore all Ag1000 sequences by default share the same indels as the PEST reference, and a tree would artifactually make all wild sequences appear more similar to one another. This analysis demonstrates that genetic variants within confirmed functional enhancer intervals, associated with differential enhancer activity, segregate in nature and do not represent variants unique to lab colonies. Natural segregation of variants associated with differential enhancer function supports the interpretation that the differential function of enhancer alleles (Figure 4) based on a modular structure of regulatory elements (Figure 6) likely result from natural selection for distinct phenotypic outcomes of allelic enhancer function.

**Figure 7.**
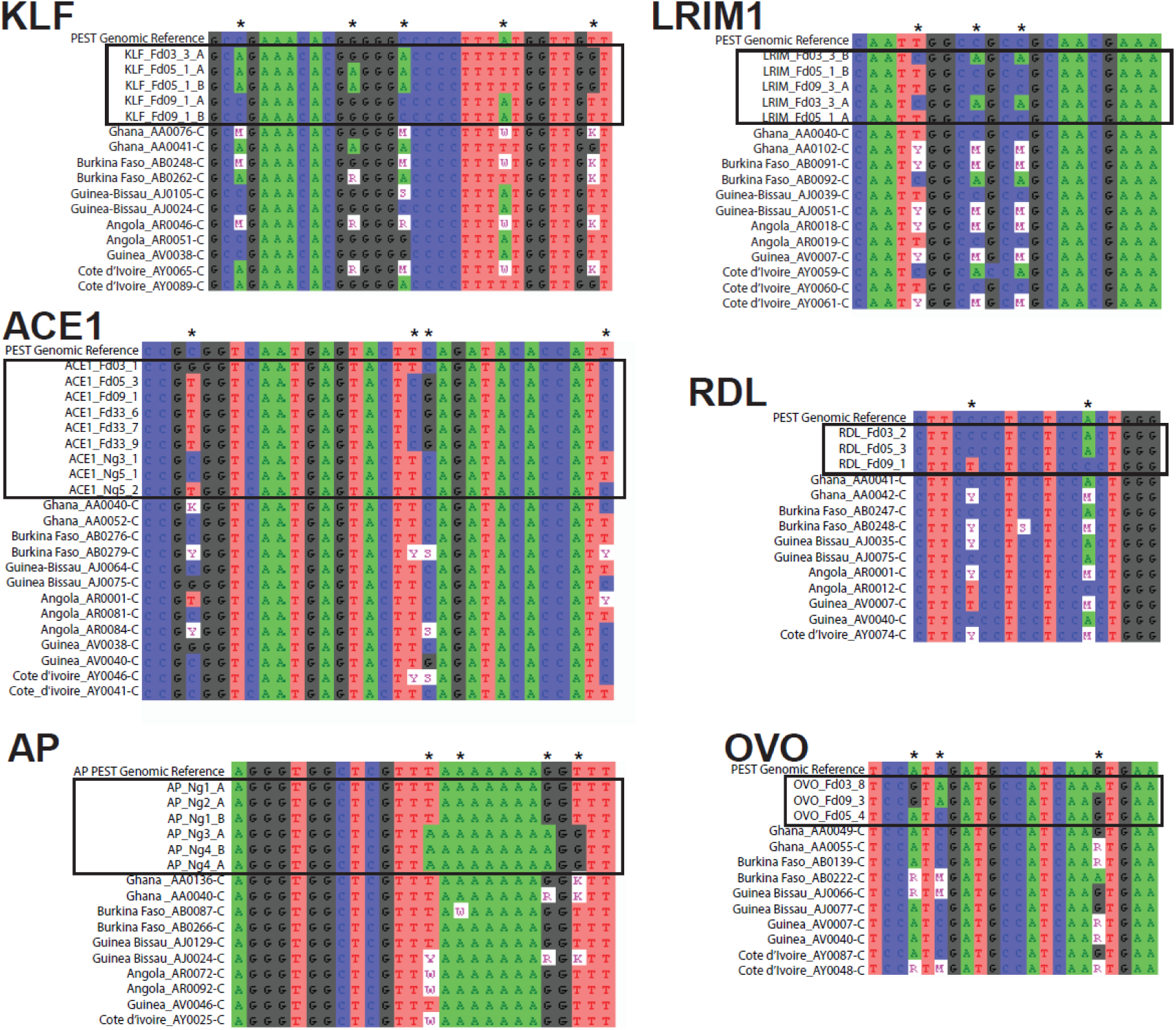
Genetic variants in differentially active enhancer alleles segregate in wild *Anopheles coluzzii*. Genetic variation observed in colonized and wild *A. coluzzii* was compared for the six studied enhancer intervals. Representative short sequence alignments are shown (full-length alignments with additional samples in Supplementary File S2). Asterisks above sequence alignments indicate variant positions shared between the cloned *A. coluzzii* colony haplotypes used in luciferase assays and the natural population. The top sequence row in each alignment is the PEST genome reference sequence, followed by sequences of alleles tested by luciferase assays (boxed by rectangles), followed by sequences of wild *A. coluzzii*. Ambiguous nucleic acid codes are used for heterozygous sites only in wild samples because the cloned sequences from *A. coluzzii* colonies are haplotypes, which are unambiguous.

## DISCUSSION

The development of a methodology for screening and evaluation of *Anopheles* enhancers is an initial step towards a more comprehensive study of enhancers and their polymorphism effects. The currently examined enhancers were chosen because they were located near known functional genes, but further work will be required to determine the actual influence, if any, of these enhancers upon the nearby genes, which is not known. Moreover, enhancer function is also controlled on spatial and temporal scales, and understanding *Anopheles* enhancers and their effects on phenotypes in detail will ultimately require incorporating this information. Lastly, the ability to translate results in the cell culture system to the whole organism will involve downstream work, but screening of enhancer candidates in a cell culture system is an important first step in determining the significance of non-coding variation to vector biology.

Here we identify and validate candidate enhancer noncoding regulatory elements in the malaria vector, *A. coluzzii*. We show that naturally segregating genetic variation significantly influences enhancer activity levels, which if replicated in vivo could result in differences in biological function and ultimately mosquito phenotype. Some enhancer alleles display high activity while others display little or no activity above background and are thus naturally occurring enhancer null alleles. The enhancers also tend to display activity that is independent of their physical orientation, a common property of enhancers ^7^. A structure-function study of two enhancer alleles by deletion mutagenesis revealed a complex modular organization of positive and negative modifiers that modulate enhancer activity. The current study provides proof of principle for the influence of enhancer genetic polymorphism for enhancer functional activity levels. These results provide benchmark parameters that can 13 now be implemented to develop a comprehensive genome-wide *Anopheles* enhancer catalog. The current work is thus a step toward the long-term goal to identify functionally important transcription factor binding motifs and correlate enhancer output with phenotypes related to *Anopheles* biology, immunity and pathogen transmission.

By modifying the level of enhancer activity, genetic variation in enhancers can cause quantitative changes in expression of the target genes regulated by the enhancers. Altered expression profiles of enhancer target genes probably in turn trigger distinct phenotypic outcomes. Different from mutations in protein coding genes, enhancers are typically located in noncoding DNA, and there is no sequence pattern to aid interpretation of noncoding variants. Here, we identified enhancer intervals proximal to genes underlying the important vector phenotypes of insecticide resistance, immunity, and development. Most mosquito studies to date have focused solely on genes and proteins associated with these traits, rather than regulatory elements controlling the genes, in part due to the limited information available for the noncoding mosquito genome.

Enhancers were identified and individual genetic variants were tested for their activity by means of standard luciferase reporter assays. Interestingly, we detected variant alleles with significant difference in their functional enhancer activity, including functional null alleles that lack enhancer activity above background. For example, the LRIM1 enhancer Fd05_#1 allele or the ACE1 Fd03_#1 allele likely represent the ablation of an important transcription factor binding site, resulting in the absence of enhancer activity above background with likely downstream functional consequences. The range in functional enhancer activity that we observed is likely to affect phenotypes produced by the genes they transcriptionally 14 regulate. Moreover, we demonstrate that variant alleles tested by luciferase activity in laboratory colonies also segregate in the wild population, and are therefore subject to natural selection. Thus, it is intriguing that selection has apparently generated a wide range of natural allelic forms of enhancers, including alleles that lack functional activity. This is consistent with the observation that genetic variation for enhancer function offers powerful raw material for adaptation and evolutionary change ^11, 20, 23, 24^. The members of the Gambiae species complex, including *A. coluzzii*, are highly adaptable to a range of ecological conditions, and durable to vector control measures. This is thought to be associated with their high genetic diversity ^40^. The current study addresses standing genetic variation and enhancer function something that needs to be considered as a likely new contributing factor influencing the success of this mosquito and its relatives.

We functionally dissected two alleles of the LRIM1 enhancer, high and low activity variants, respectively, by deletion mutagenesis (Figure 6). By measuring functional activity of integral, 50% and 25% length derivatives of the intervals, we detected a modular structure of positive and negative regulators comprising the enhancer. Interestingly, deletion derivatives of the two alleles behaved differently, indicating that the deletions was not a simple consequence of sequence length. The high-activity allele Fd03_#3 appears to carries a negative regulator in the terminal one-quarter of its length on one or both ends, because removal of these sequences led to significantly elevated activity in the remaining 50% derivative as compared to the integral enhancer. However, removal of an additional one-quarter again of the sequence from both ends of the 50% derivative then diminished activity to a level below that of the integral enhancer, suggesting that the positive regulator(s) revealed in the 50% length derivative was no longer present in the 25% length derivative.

That the low activity of the smallest derivative of Fd05_#1 was not a simple consequence of sequence length is made clear by a similar examination of the low-activity allele Fd05_#1. In this case, each incremental length decrease of the tested sequence led to increased enhancer activity. The Fd05_#1 allele result suggests that the integral enhancer displayed low activity because it carried multiple negative regulators, which were resected by each successive deletion, revealing a highly active core enhancer element within the smallest interval tested. This latter minimal derivative of the low-activity Fd05_#1 allele carries an enhancer with, in fact, higher enhancer activity than the integral 100% sequence of the high-activity Fd03_#3 allele.

The LRIM1 deletion mutagenesis results suggest that that large functional allelic diversity can be generated for a given enhancer interval by the combinatorial effect of positive and negative modifiers. Sequence changes in enhancers can generate or remove binding motifs for transcription factors and other regulatory proteins, which can modify transcription levels directly ^41, 42^, or indirectly through loss of chromatin accessibility ^17^ From the current results, we do not know whether the positive and negative modifiers within the LRIM1 alleles are comprised of reusable modular cassettes that are combined to fine-tune the activity of different enhancer intervals, or whether segregating SNPs in an enhancer can explain significant difference in functional activity. In the first model, such modifier modules should be recognizable with enough data, while under the second model, different modifiers, for example positive modifiers in different enhancers, may have little or no recognizable common pattern. Fine resolution nucleotide-level deletion series of a number of alleles would be required to determine the kind and extent of sequence difference necessary to 16 alter enhancer functional levels in order to distinguish between the above models. Finally, the phenotypic implications of differentially active enhancer alleles will require determination of target gene networks regulated by an enhancer, as well as the protein-DNA interactions underlying differential enhancer allele activity.

Vector control has been central to the malaria control effort by use of indoor residual spraying and long-lasting insecticide impregnated bednets. However, over-reliance on these methods has led to widespread insecticide resistance in wild populations, and novel methods of control are now required. The noncoding regulatory genome in *Anopheles* has the potential to provide novel new targets for vector control, but until now has not been interpretable or exploitable. For example, understanding the regulatory genome landscape could improve the choice of insertion sites for exogenous transgenes for proper expression, and enhancers themselves could be targeted in order to alter transcription of immune or disease important genes. The current work presents a necessary first step towards establishing an efficient, effective method for associating noncoding variation with important mosquito phenotypes.

## METHODS

### Wild mosquito samples and DNA library

Mosquito larvae were collected in Goundry village, Burkina Faso (latitude 12.5166876, longitude −1.3921092) using described methods ^43^, reared to adults, and were typed for species by the SINE200 X6.1 assay ^44^. DNA from 60 *A. coluzzii* were pooled at equal volume and sheared using an S220 ultrasonicator (Covaris) to produce DNA fragments 800-1000 bp in length. Subsequently, DNA was processed as described for the STARR-seq assay ^35^, cloned into the plasmid pSTARR-seq_fly (AddGene 71499), transformed into MegaX DH10B T1R Electrocomp Cells (Invitrogen), cultured in LB + ampicillin (1ug/ml), and plasmid DNA was purified using the Plasmid Plus Mega Kit (Qiagen). The *Anopheles* gambiae PEST AgamP4 genome assembly available at Vectorbase was used as the reference genome (https://www.vectorbase.org/organisms/anopheles-gambiae/pest/agamp4).

### Culture of plasmid library in *Anopheles* 4a3A cells

Hemocyte-like 4a3A cells ^45^ were maintained on Insect X-Press media (Lonza) supplemented with 10% Fetal Calf Serum (heat inactivated at 56°C for 30 minutes), at 27°C. No antibiotics were used. We confirmed that cells were derived from *A. coluzzii* by species typing using the Fanello assay ^46^. The plasmid DNA library was transfected and cultured in 4a3A cells as described ^35^ using Lipofectamine 3000 Reagent (Invitrogen) and cultured for 24 h, in three biological replicates. RNA was extracted from cells using the RNeasy Midi Kit (Qiagen) followed by mRNA purification using Dynabeads mRNA Purification Kit (ThermoFisher). Plasmid DNA was isolated using the Plasmid Plus Midi or Mini Kit (Qiagen).

### Analysis of 4a3A library culture results

The mRNA purified from cells was reverse transcribed using SuperScript IV First-Strand cDNA Synthesis System (Invitrogen) as described for the STARR-seq assay ^35^ using a plasmid-specific primer (RT_Rev, Supplementary Table S1), the cDNA was then amplified using primers Report_Fwd and Report_Rev (Supplementary Table S1), and the products were sequenced on an Illumina HiSeq 2500 in 2×125 bp high output mode. Cell plasmid DNA was amplified and sequenced in the same manner as the cDNA samples but using primers Plasmid_Fwd and Plasmid_Rev (Supplementary Table S1).

### Selection of enhancer candidates

The Integrative Genomics Viewer (IGV) ^36^ was used to select candidate enhancer intervals by visual examination in the proximity of six annotated genes of interest. For determination of enhancer activity, the RNA output transcribed from the STARR-seq reporter plasmid, converted to cDNA and sequenced as described above, is compared to the levels of the plasmid DNA, to control for differential plasmid replication. Thus, candidate enhancers were predicted in intervals where coverage of the cDNA sequence track was visibly greater than the baseline coverage of the plasmid sequence track. The target genes examined were Krueppel-Like Factor 6/7 (KLF, AGAP007038), Leucine-Rich Immune protein (LRIM1, AGAP006348), Acetylcholinesterase (ACE1, AGAP001356), GABA-gated chloride channel subunit (Rdl, AGAP006028), LIM homeobox protein 2/9, ortholog of Drosophila apterous FBgn0267978 (AP, AGAP008980), and Ovo, AGAP000114. In addition, a negative control interval was cloned, which was a size-matched interval located within intron 1 of the gene, homeobox protein distal-less (DLX, AGAP007058) that displayed no visible divergence of cDNA and plasmid sequence tracks by IGV examination, and thus no predicted enhancer function. In all but one case, the candidate enhancer was the only one in the vicinity of the 19 target gene, for AP there were three peaks, the most gene proximal peak is likely a promoter so the next most proximal peak was chosen as shown in Figure 1. The candidate enhancer intervals are named according to the most proximal coding sequence (above and Table 1).

Candidate enhancers were amplified from DNA of mosquitoes from the following *A. coluzzii* colonies: Ngousso, initiated in Cameroon in 2006 ^47^, Fd03, Mali, 2008, Fd05, Mali 2008, Fd09, Burkina Faso, 2008, and Fd33, Burkina Faso, 2014. Fd colonies were previously described ^48^. Primers are listed in Supplementary Table S2. Amplicons were cloned into the pCR8/GW/TOPO vector (Invitrogen) and sequenced with GW1 and GW2 primers. A standard plasmid was used for luciferase assays that incorporates the engineered Drosophila synthetic core promoter (DSCP). The information about the fine structure of Anopheles core promoters does not yet exist. The results indicate that the DSCP is functional in Anopheles cells. At least two genetically distinct sequences per candidate were then cloned into the firefly luciferase reporter plasmid pGL-Gateway-DSCP (AddGene 71506) using Gateway LR Clonase II (Invitrogen), transformed into OneShot OmniMax 2T1 Phage-Resistant Cells (Invitrogen), and plasmid was purified from overnight culture.

To test the effect of enhancer orientation, the enhancer was cloned in the opposite orientation in pGL-Gateway-DSCP. To test resolved enhancer intervals, the relevant enhancer insert was amplified with primers that generated either 50% or 25% of the initial insert size, equally reduced on both ends, and products were cloned in pGL-Gateway-DSCP. In all cases, plasmids were resequenced to confirm insert identity using the primers LucNrev and RVprimer3 (Supplementary Table S1).

### Quantitation and statistical analysis of enhancer activity by luciferase assay

The Dual-Glo Luciferase Assay System (Promega) was used for luciferase assays. *A. coluzzii* 4a3A cells were seeded in 96 well plates at 1×10^5^cells/well, the difference in volume if any was made up to 65 μl with medium, and cells were incubated for 24 h at 27°C. Two plasmids were transfected into the 4a3A cells, the enhancer candidate in firefly luciferase vector pGL-Gateway-DSCP, and the renilla luciferase control vector pRL-ubi-63E (AddGene 74280), at a ratio of 1:5 (renilla:firefly), using transfection reagent Lipofectamine 3000 (Invitrogen), and were then incubated for 24 h at 27°C.

Luciferase activity was detected on a GloMax Discover instrument (Promega) at 25°C, with two 20 min incubations, one after the addition of Dual-Glo Luciferase reagent (Promega) and another after the addition of Stop & Glo reagent (Promega). All samples were run in 6-fold replication within a single plate and across at least two independent plates, for at least two biological replicates of each candidate, yielding at least 12 measurements. Firefly luciferase measurements expressed in relative light units (RLU) were corrected using the measurements of RLU for the renilla luciferase internal control in the same well. Values for the DLX negative control on the same plate were defined as the background level. Values of firefly/renilla RLU for the experimental insert were normalized to the firefly/renilla mean value for DLX in order to combine results across replicates. Luciferase activity was declared above background if the firefly/renilla RLU ratio for the experimental insert was significantly higher than the firefly/renilla value for the DLX negative control. Luciferase activity was statistically compared using a non-parametric ANOVA (Kruskal-Wallis) with post hoc pairwise comparisons.

### Analysis of enhancer allelic variants

The sequences of genetically polymorphic variants of a given enhancer interval, cloned from *A. coluzzii* colonies as described above, were analyzed for genetic relatedness. To generate neighbor joining (N-J) trees to depict the relationships between genetic variants for the same enhancer, complete sequences were aligned using MUSCLE within the package Molecular Evolutionary Genetics Analysis Mega version X ^49^, and N-J trees constructed using Mega. When at least four variants were tested, bootstrapping was performed and bootstrap values are included on N-J trees. Scale bars of trees represent 0.5, but each bar is a different length. The longer the 0.5 scale bar, the more genetically similar the sequences.

### Analysis of wild *Anopheles* variation data

Sequence information for 309 wild *A. coluzzii* from 6 West African countries; Angola (AR), Burkina Faso (AB), Cote d’Ivoire (AY), Ghana (AA), Guinea (AV) and Guinea-Bissau (AJ), generated as reported ^40^ were downloaded from MalariaGen (https://www.malariagen.net/projects/ag1000g) as VCF files from the Ag1000G phase 2 AR1 data release and sequences of the six validated enhancer intervals were extracted from the raw VCF files using GATK version 3.9 (SelectVariants mode) ^50^, the consensus sequences with IUPAC ambiguity codes for the variants were extracted using BCFtools version 1.9 (consensus mode with –iupac-codes option) ^51^, the sequences of each interval were aligned with Clustal W version 2.1 ^52^ and visualized with AliView aligner version 1.24 ^53^.

The diploid sequences from the wild sequences were aligned to the cloned sequences generated from the six validated enhancer intervals. Sequence alignments were visually examined for shared variation. Short representative sequence alignments are presented in 22 Figure 7 (not including indels), and complete alignments relative to the PEST AgamP3genome assembly, including indels, are presented in Supplementary File S2. Indel genotypes of the wild sequences shown in Supplementary File S2 are relative to the PEST reference haplotype, because the wild sequences were called for SNPs but not indels ^40^.

## Supporting information

Supplementary Figure 1 and other Supplementary Information

## Acknowledgments

We thank the Center for Production and Infection of *Anopheles* platform (CEPIA) at the Institut Pasteur, and Corinne Genève and Emma Brito-Fravallo, GGIV Institut Pasteur, for rearing mosquitoes. We thank Alexander Stark, Research Institute of Molecular Pathology, Vienna for plasmids and helpful advice. This work received financial support to MMR from National Institutes of Health, NIAID #AI121587; to KDV from the European Commission, Horizon 2020 Infrastructures #731060 Infravec2; European Research Council, Support for Frontier Research, Advanced Grant #323173 AnoPath; and French Laboratoire d’Excellence “Integrative Biology of Emerging Infectious Diseases” #ANR-10-LABX-62-IBEID. The funders had no role in study design, data collection and analysis, decision to publish, or preparation of the manuscript.

## Author contributions statement

Conceived and designed the experiments: MMR, DMG, KDV

Performed the experiments: LN, IH, DMG, SZ, WMG, NS, MMR

Analyzed the data: AP, EB, MMR

Wrote the manuscript: LN, MMR, KDV

All authors read and approved the final manuscript.

## Ethics statement

No animals or human subjects were used. Mosquito colonies were maintained on anonymous commercial human blood using an artificial membrane feeding device.

## Data availability

All short read sequence files are available from the EBI European Nucleotide Archive database (http://www.ebi.ac.uk/ena/) under ENA study accession number [REQUESTED]. All other sequences are available in this article as Supplementary Files S1 and S2.

## Competing interests statement

The authors declare no competing interests.

## Notes

#### Summary of Updates

This version of the manuscript is updated for greater clarity, more experimental details and a workflow summary figure to help better understand the workflow that generated the presented data. We also make clear the power and limits of the approach used.

