## Supplementary Figure 1 and other Supplementary Information for "Influence of genetic polymorphism on transcriptional enhancer activity in the malaria vector *Anopheles coluzzii*"

**Supplementary Information Legends**

**Supplementary Table S1. Primers used in DNA cloning and cDNA sequence analysis.** List of all primers used for Anopheles coluzzii library cloning and analysis of cDNA after growth in A. coluzzii 4a3A cells.

**Supplementary Table S2. Primers used to amplify Anopheles coluzzii candidate enhancer intervals.** List of all primers targeting A. coluzzii genomic sequence.

**Supplementary Figure S1. Experimental work flow.** **1**. Enrichment of transcript sequence reads above control reads identifies enhancer fragments near the immune, insecticide and development genes of interest. **2.** Use mosquito genomic DNA and primers flanking the enhancer region to amplify the enhancer region of interest (in blue, not to scale). **3.** Clone the enhancer allele (blue) into an expression vector where it controls expression of the luciferase gene (yellow arrow) and light output from a luciferase assay is proportional to enhancer activity. The greater the light activity is, the more enhancer activity the candidate enhancer fragment (blue) has. **4.** Transfect the luciferase reporter construct from step 3 into mosquito 4a3a cells and measure light output with a luminometer. Normalize all luciferase activity to light output from a size matched genomic fragment that does not show enrichment of transcript sequence (i.e. showed no peak in step 1). **5.** Amplify the enhancer fragment from a number of mosquito individuals to identify genetic variants and clone these variants (represented as colored bars on the blue background) into the expression vector (step 3) to measure the effects of genetic variation on enhancer activity as determined by luciferase activity and light output.

**Supplementary File S1. Complete genomic sequences for all tested Anopheles coluzzii enhancer alleles**. Complete information for sequences of N-J tree in Figure 3. Amplicons generated using primers presented in Supplementary Table S1. Each file contains aligned allele sequences for each enhancer.

**Supplementary File S2. Alignments of tested Anopheles coluzzii enhancer intervals sequenced from laboratory colonies and wild population individuals**. Full-length alignments corresponding to short sequence windows presented in Figure 7, with larger numbers of wild mosquito sequences.

**Supplementary Table S1. Primers used in DNA cloning and cDNA sequence analysis.**

| **Primer name** | **Sequence (5’-3’)** |
| --- | --- |
| STARR_Seq_PCR1_For | TAG AGC ATG CAC CGG ACA CTC TTT CCC TAC ACG ACG CTC TTC CGA TCT |
| STARR_Seq_PCR1_Rev | GGC CGA ATT CGT CGA CAA GCA GAA GAC GGC ATA CGA GAT |
| RT_Rev | CAA ACT CAT CAA TGT ATC TTA TCA TG |
| Report_Fwd | AAG CCA CCA TGG AAA AGG CCA T |
| Report_Rev | TAT CAT GTC TGC TCG AAG CGG |
| Plasmid_Fwd | CTA GGC ACA CCG AAA CGA CTA AC |
| Plasmid_Rev | TAT CAT GTC TGC TCG AAG CGG |
| D501_TS_primer | AAT GAT ACG GCG ACC ACC GAG ATC TAC ACT ATA GCC TAC ACT CTT TCC CTA CAC GAC GCT CTT CCG ATC T |
| D502_TS_primer | AAT GAT ACG GCG ACC ACC GAG ATC TAC ACA TAG AGG CAC ACT CTT TCC CTA CAC GAC GCT CTT CCG ATC T |

**Supplementary Table S2. Primers used to amplify Anopheles coluzzii candidate enhancer intervals**. Fwd, forward primer, Rev, reverse primer, bp, base pairs.

| **Primer name** | **AGAP ID** | **Amplicon size (bp)** | **Primer sequence (5’-3’)** |
| --- | --- | --- | --- |
| AP Fwd | AGAP008980 | 1200 | TTGTGCATCGCTTGAAAGAA |
| AP Rev |  |  | TTCATGAGAAGAATCAAAGAGACAA |
| OVO Fwd | AGAP000114 | 1551 | TGGCACTCCAAAGTGTAGGG |
| OVO Rev |  |  | TCTTCCACCGCTTCAAGTTC |
| KLF Fwd | AGAP007038 | 1240 | TTGCATACCTTCAGGCGTTT |
| KLF Rev |  |  | ACCATGGATGCCCCTAAAAG |
| LRIM1 Fwd | AGAP006348 | 1357 | CTTTCCCCTCTTGCGTGTAG |
| LRIM1 Rev  LRIM1 Left 50%  LRIM1 Right 50%  LRIM1 Left 25%  LRIM1 Right 25% |  | 737  396 | AGACGATACAGCAGCAGCAA  GCTTGAGCGCTGTCTGCCCTC  GTAAGAAGTTGCTTTTACCCAGA  CAGCCTGCGTACCGGCAATG  CAGCGATGCGAAACGGGTCTGC |
| ACE Fwd | AGAP001356 | 1148 | CCTTTCTTCCAGCCACCTTT |
| ACE Rev |  |  | TTGTGCTCGGATTGCTATGA |
| RDL Fwd | AGAP006028 | 1426 | TCGTTCAGCGTTGGTTATGA |
| RDL Rev |  |  | TACGCTTGTTCGCACACTTT |
| DLX Fwd | AGAP007058 | 960 | GCTTACGCCATCTGGTTGAT |
| DLX Neg |  |  | CAGTGGTGTTTTGCATTTCG |

**Supplementary Figure S1. Experimental work flow**

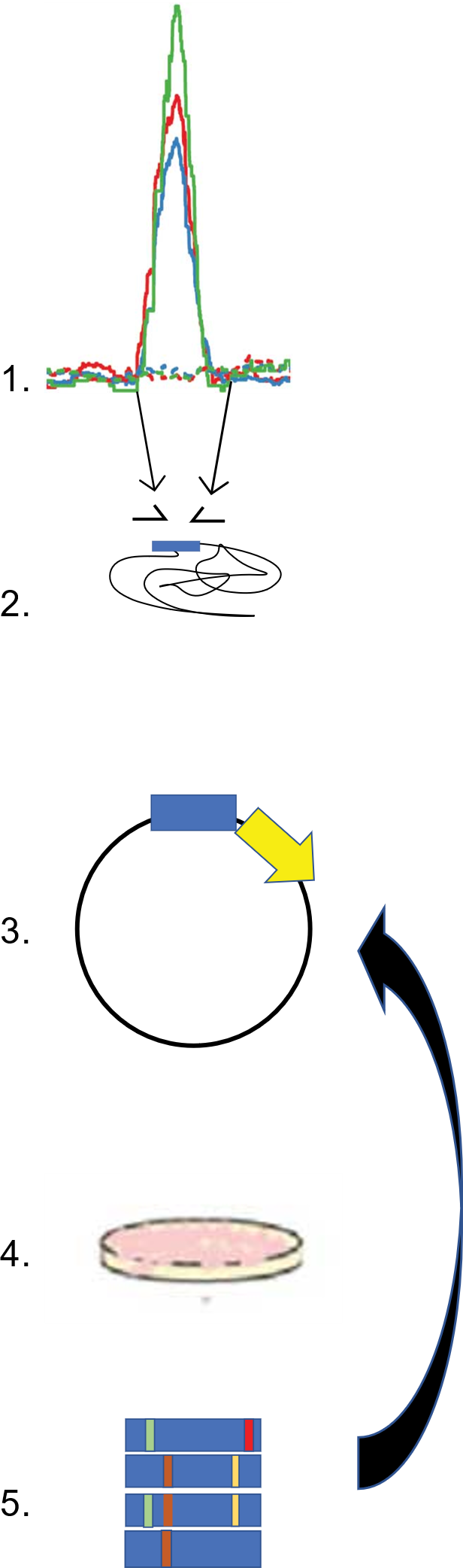
